# Whole genome analysis illustrates global clonal population structure of the ubiquitous dermatophyte pathogen *Trichophyton rubrum*

**DOI:** 10.1101/261743

**Authors:** Gabriela F. Persinoti, Diego A. Martinez, Wenjun Li, Aylin Döğen, R. Blake Billmyre, Anna Averette, Jonathan M. Goldberg, Terrance Shea, Sarah Young, Qiandong Zeng, Brian G. Oliver, Richard Barton, Banu Metin, Süleyha Hilmioğlu-Polat, Macit Ilkit, Yvonne Gräser, Nilce M. Martinez-Rossi, Theodore C. White, Joseph Heitman, Christina A. Cuomo

**Author notes:** equal contribution as first authors (Gabriela F. Persinoti, Diego A. Martinez, Wenjun Li). Current addresses: Veritas Genetics, Danvers, Massachusetts 01923, USA. National Center for Biotechnology Information (NCBI), 8600 Rockville Pike, Bethesda, MD 20894, USA. Stowers Institute for Medical Research, Kansas City, Missouri, USA. Harvard T. H. Chan School of Public Health, Boston, Massachusetts 02115, USA. LabCorp, Westborough, MA 01581, USA. corresponding authors.Corresponding authors: Christina A. Cuomo, Broad Institute, 415 Main Street, Cambridge, MA 02142. Joseph Heitman, 322 CARL Building, Box 3546, Duke University Medical Center, Durham, N.C. 27710.

## Abstract

Dermatophytes include fungal species that infect humans, as well as those which also infect other animals or only grow in the environment. The dermatophyte species *Trichophyton rubrum* is a frequent cause of skin infection in immunocompetent individuals. While members of the *T. rubrum* species complex have been further categorized based on various morphologies, the population structure and ability to undergo sexual reproduction are not well understood. In this study, we analyze a large set of *T. rubrum* and *Trichophyton interdigitale* isolates to examine mating types, evidence of mating, and genetic variation. We find that nearly all isolates of *T. rubrum* are of a single mating type, and that incubation with *T. rubrum* morphotype *megninii* isolates of the other mating type failed to induce sexual development. While the region around the mating type locus is characterized by a higher frequency of SNPs compared to other genomic regions, we find that the population is remarkably clonal, with highly conserved gene content, low levels of variation, and little evidence of recombination These results support a model of recent transition to asexual growth when this species specialized to growth on human hosts.

Data access: Genome sequence data is available in NCBI under the Umbrella BioProject PRJNA186851.

## Introduction

Dermatophyte species are the most common fungal species causing skin infections. Of the more than 40 different species infecting humans, *Trichophyton rubrum*, the major cause of athlete’s foot, is the most frequently observed [1,2]. Other species are more often found on other skin sites, such as those found on the head, including *Trichophyton tonsurans* and *Microsporum canis*. Some dermatophyte species only cause human infections, including *T. rubrum*, *T. tonsurans*, and *T. interdigitale*. Other species, including *Trichophyton benhamiae, Trichophyton equinum*, *Trichophyton verrucosum*, and *M. canis*, infect mainly animals and occasionally humans, while others such as *Microsporum gypseum* ( *Nannizzia gypsea* [3]) are commonly found in soil and rarely infect animals. In addition to the genera *Trichophyton* and *Microsporum*, *Epidermophyton* and *Nannizzia* are other genera of dermatophytes that commonly cause infections in humans [4]. The species within these genera are closely related phylogenetically and are within the Ascomycete order Onygenales, family Arthrodermatacaea [3,5].

The *Trichophyton rubrum* species complex includes several “morphotypes,” many of which rarely cause disease, and *T. violaceum*, a species that causes scalp infections [3,6]. Some morphotypes display phenotypic variation, though these differences can be modest. For example, *T. rubrum* morphotype *raubitscheckii* differs from *T. rubrum* in production of urease and in colony pigmentation and colony appearance under some conditions [7]. *T. rubrum* morphotype *megninii*, which is commonly isolated in Mediterranean countries, requires L-histidine for growth unlike other *T. rubrum* isolates [6]. However, little variation has been observed between these and other morphotypes in the sequence of individual loci, such as the ITS rDNA locus; additionally, some of the morphotypes do not appear to be monophyletic [3,6,8], complicating any simple designation of all types as separate species. Combining morphological and multilocus sequence typing (MLST) data has helped clarify relationships of the major genera of dermatophytes and resolved polyphyletic genera initially assigned by morphological or phenotypic data.

Mating has been observed in some dermatophyte species, although not to date in strict anthropophiles including *T. rubrum* [9]. Mating type in dermatophytes, as in other Ascomycetes, is specified by the presence of one of two idiomorphs at a single mating type (*MAT*) locus; each idiomorph includes either an alpha box domain or HMG domain transcription factor gene [10]. In the geophilic species *M. gypseum*, isolates of opposite mating type ( *MAT1-1* and *MAT1-2*) undergo mating and produce recombinant progeny [10]. In the zoophilic species *T. benhamiae*, both mating types are detected in the population and mating assays produced fertile cleistothecia [11], structures that contain meiotic ascospores. In a study examining 600 isolates of *T. rubrum*, only five appeared to produce structures similar to cleistothecia [12], suggesting inefficient development of the spores required for mating. Sexual reproduction experiments of *T. rubrum* with tester strains of *Trichophyton simii*, a skin infecting species that is closely related to *T. mentagrophytes*, have been reported and one recombinant isolate was characterized, consistent with a low frequency of mating of *T. rubrum* [13]. Further, sexual reproduction of *T. rubrum* may be rare in natural populations, as a single mating type (*MAT1-1*) has been noted in Japanese isolates [14], matching that described in the *T. rubrum* reference genome of CBS 118892 [10].

Here we describe genome-wide patterns of variation in *T. rubrum*, revealing a largely clonal population. This builds on prior work to produce reference genomes for *T. rubrum* [15] and other dermatophytes [15,16]. Genomic analysis of two divergent morphotypes of *T. rubrum*, *megninii* and *soudanense*, reveal hotspots of variation linked to the mating type locus suggestive of recent recombination. While nearly all *T. rubrum* isolates are of a single mating type (*MAT1-1*), the sequenced *megninii* morphotype isolate contains a *MAT1-2* locus, suggesting the capacity for infrequent mating in the population. Additionally, we examine variation in gene content across dermatophyte genomes including the first representatives of *T. interdigitale*.

## Materials and methods

### Isolate selection, growth conditions, and DNA isolation

Isolates analyzed are listed in **Table S1**, including the geographic origin, site of origin, and mating type for each. Isolates selected for whole genome sequencing were chosen to maximize diversity by covering the main known groups. For whole genome sequencing, 10 *T. rubrum* isolates and 2 *T. interdigitale* isolates were selected, including representatives of the major morphotypes of *T. rubrum* (**Table S2**). Growth and DNA isolation for whole genome sequencing were performed as previously described [15].

For MLST analysis, a total of 80 *T. rubrum* isolates and 11 *T. interdigitale* isolates were selected for targeted sequencing. Isolates were first grown on PDA medium (Difco) for 10 days at 25°C. Genomic DNA was extracted using an Epicentre Masterpure Yeast DNA purification kit (catalog number MPY08200). Fungal isolates were harvested from solid medium using sterile cotton swabs, transferred to microcentrifuge tubes, and washed with sterile PBS. Glass beads (2 mm) and 300 μL yeast cell lysis solution (Epicentre) were added to the tube to break down fungal cells, and the protocol provided by Epicentre was then followed. The contents of the tube were mixed by vortexing and incubated at 65**°**C for 30 minutes, followed by addition of 150 μl Epicentre MPC Protein Precipitation Solution. After vortexing, the mixture was centrifuged for 10 minutes, followed by isopropanol precipitation and washing with 70% ethanol. The DNA pellet was dissolved in TE buffer.

For mating assays, we investigated 55 *T*. *rubrum* and 9 *T*. *interdigitale* isolates recovered from Adana and Izmir, Turkey. *T. simii* isolates CBS 417.65 MT -, CBS 448.65 MT + and morphotype *megninii* isolates CBS 389.58, CBS 384.64, and CBS 417.52 were also used in mating assays. DNA extraction was performed according to the protocol described by Turin et al. [17]. These isolates were typed by ITS sequence analysis. rDNA sequences spanning the internal transcribed spacer (ITS) 1 region were PCR-amplified using the universal fungal primers ITS1 (5’-TCCGTAGGTGAACCTGCGG3’) and ITS4 (5’-CCTCCGCTTATTGATATGC-3’) and sequenced on an ABI PRISM 3130XL genetic analyzer at Refgen Biotechnologies using the same primers (Ankara, Turkey). CAP contig assembly software, included in the BioEdit Sequence Alignment Editor 7.0.9.0 software package, was used to edit the sequences [18]. Assembled DNA sequences were characterized using BLAST in GenBank.

### Multilocus sequence typing (MLST)

A total of 108 isolates were subjected to MLST analysis (**Table S3**). For each isolate, three loci (the *TruMDR1* ABC transporter [19], an intergenic region (IR), and an alpha-1,3-mannosyltransferase (CAP59 protein domain)), with high sequence diversity between *T. rubrum* CBS 118892 (GenBank accession: NZ_ACPH00000000) and *T. tonsurans* CBS 112818 (GenBank accession: ACPI00000000), were selected as molecular markers in MLST. The following conditions were used in the PCR amplification of the three loci: an initial 2 min of denaturation at 98°C, followed by 35 cycles of denaturation for 10 sec at 98°C, an annealing time of 15 sec at 54°C, and an extension cycle for 1 min at 72°C. The amplification was completed with an extension period of 5 min at 72°C. PCR amplicons were sequenced using the same PCR primers on an ABI PRISM 3130XL genetic analyzer by Genewiz, Inc. (**Table S4**). Electropherograms of Sanger sequencing were examined and assembled using Sequencher 4.8 (Gene Codes). Alternatively, sequences were obtained from genome assemblies (**Table S5**).

To confirm the species typing for four isolates (MR857, MR827, MR816, and MR897), the ITS1, 5.8S, and ITS2 region was amplified using the ITS5 (5’-GAAGTAAAAGTCGTAACAAGG-3’) and Mas266 (5’-GCATTCCCAAACAACTCGACTC-3’) primers with initial denaturation at 94°C for 4 minutes, 35 cycles of denaturation at 94°C for 30 seconds, annealing at 60°C for 30 seconds, extension at 72°C for 1 minute, and final extension at 72°C for 10 minutes. The reactions were carried out using a BioRad C1000 Touch thermocycler. ABI sequencing reads were compared to the dermatophyte database of the Westerdijk Fungal Biodiversity Institute. The sequences of MR857 and MR827 isolates were 99.6% identical to that of the isolate RV 30000 of the African race of *T. benhamiae* (GenBank AF170456).

### Mating type determination

To identify the mating type of each isolate, primers were designed to amplify either the alpha or HMG domain of *T. rubrum* (**Table S6**). For most isolates, PCR amplification was performed using an Eppendorf epGradient Mastercycler, and reactions were carried out using the following conditions for amplification: initial denaturation at 94°C for 4 minutes, 35 cycles of denaturation at 94°C for 30 seconds, annealing at 55°C for 30 seconds, extension at 72°C for 1 minute, with a final extension at 72°C for 7 minutes. For isolates from Turkey, PCR amplifications were performed with the same primers using a Biorad C1000 Touch^TM^ Thermal Cycler, and slightly modified conditions were used for amplification: initial denaturation at 94oC for 5 minutes, 35 cycles of denaturation at 95oC for 45 seconds, annealing at 55oC for 1.5 minutes, and extension at 72oC for 1 minute, and final extension at 72oC for 10 minutes. The presence of the alpha box gene, which is indicative of the *MAT1-1* mating type, or the HMG domain, which is indicative of the *MAT1-2* mating type, was identified using primers JOHE21771/WL and JOHE21772/WL, creating a 500-bp product, and JOHE21773/WL and JOHE21774/WL, creating a 673-bp product, respectively. *Trichophyton rubrum* MR 851 was used as a positive control for *MAT1-1*, and morphotype *megninii* CBS 389.58, CBS 384.64, CBS 417.52, and *T. interdigitale* MR 8801 were used as positive controls for *MAT1-2*. The mating type was assigned based on the presence or absence of PCR products on 1.5% agarose gels. For the whole genome sequenced isolates, mating type was determined by analysis of assembled and annotated genes.

### Mating assays

Mating assays were performed using both Medium E (12 g/L oatmeal agar (Difco), 1 g/L MgSO_4_.7H_2_O, 1 g/L NaOH_3_, 1g/L KH_2_PO_4_, and 16 g/L agar [20]) and Takashio medium (1/10 Sabouraud containing 0.1% neopeptone, 0.2% dextrose, 0.1% MgSO_4_.7H_2_O, and 0.1%. KH_2_PO_4_). *MAT1-1* and *MAT1-2* isolates grown on Sabouraud Dextrose Agar (SDA) for one week were used to inoculate both Medium E and Takashio medium plates pairwise 1 cm apart from each other. The plates were incubated at room temperature without Parafilm in the dark for 4 weeks. The petri dishes were examined under light microscopy for sexual structures.

### Genome sequencing, assembly, and annotation

For genome sequencing, we constructed a 180-base fragment library from each sample, by shearing 100 ng of genomic DNA to a median size of ∼250 bp using a Covaris LE instrument and preparing the resulting fragments for sequencing as previously described [21]. Each library was sequenced each on the Illumina HiSeq 2000 platform. Roughly 100X of 101 base-paired Illumina reads were assembled using ALLPATHS-LG [22] run with assisting mode utilizing *T. rubrum* CBS118892 as a reference. For most genomes, assisting mode 2 was used (ASSISTED_PATCHING=2) with version R42874; for *T. interdigitale* H6 and *T. rubrum* MR1463 version R44224 was used. For *T. rubrum* morphotype *megninii* CBS 735.88 and *T. rubrum* morphotype *raubitschekii* CBS 202.88 mode 2.1 was used (ASSISTED_PATCHING=2.1) with version R47300. Assemblies were evaluated using GAEMR (http://software.broadinstitute.org/software/gaemr/); contigs corresponding to the mitochondrial genome or contaminating sequence from other species were removed from assemblies.

The *Trichophyton* assemblies were annotated using a combination of expression data, conservation information, and *ab-initio* gene finding methods as previously described [23]. Expression data included Illumina reads (SRX123796) from one RNA-Seq study [24] and all EST data available in GenBank as of 2012. RNA-Seq reads were assembled into transcripts using Trinity [25]. PASA [26] was used to align the assembled transcripts and ESTs to the genome and identify open reading frames (ORFs); gene structures were also updated in the previously annotated *T. rubrum* CBS118892 assembly [15]. Conserved loci were identified by comparing the genome with the UniRef90 database [27] (updated in 2012) using BLAST [28]. The BLAST alignments were used to generate gene models using Genewise [29]. The *T. rubrum* CBS 118892 genome was aligned with the new genomes using NUCmer [30]. These alignments were used to map gene models from *T. rubrum* to conserved loci in the new genomes.

To predict gene structures, GeneMark, which is self-training, was applied first; GeneMark models matching GeneWise ORF predictions were used to train the other *ab-initio* programs. *Ab-intio* gene-finding methods included GeneMark [31], Augustus [32], SNAP [33] and Glimmer [34]. Next, EVM [35] was used to select the optimal gene model at each locus. The input for EVM included aligned transcripts from Trinity and ESTs, gene models created by PASA and GeneWise, mapped gene models, and *ab-initio* predictions. Rarely, EVM failed to produce a gene model at a locus likely to encode a gene. If alternative gene models existed at such loci, they were added to the gene set if they encoded proteins longer than 100 amino acids, or if the gene model was validated by the presence of a PFAM domain or expression evidence. Finally, PASA was run again to improve gene-model structure, predict splice variants, and add UTR.

Gene model predictions in repetitive elements were identified and removed from gene sets if they overlapped TPSI predictions (http://transposonpsi.sourceforge.net), contained PFAM domains known to occur in repetitive elements, or had BLAST hits against the Repbase database [36]. Additional repeats were identified using a BLAT [37] self-alignment of the gene set to the genomic sequence (requiring at least 90% nucleotide identity over 100 bases aligned); genes that hit the genome more than eight times using these criteria were removed. Genes with PFAM domains not found in repetitive elements were retained in the gene set, even if they met the above criteria for removing likely repetitive elements from the gene set.

Lastly, the gene set was inspected to address systematic errors. Gene models were corrected if they contained in-frame stop codons, had coding sequence overlaps with coding regions of other gene models or predicted transfer or ribosomal RNAs, contained exons spanning sequence gaps, had incomplete codons, or with UTRs overlapping the coding sequences of other genes. Transfer RNAs were predicted using tRNAscan [38], and ribosomal RNAs were predicted with RNAmmer [39].

All annotated assemblies and raw sequence reads are available in NCBI (**Table S5**).

### SNP identification and classification

To identify SNPs within the *T. rubrum* group, Illumina reads for each *T. rubrum* isolate were aligned to the *T. rubrum* CBS 118829 reference assembly using BWA-MEM [40]; reads from the H6 *T. interdigitale* were also aligned to the *T. interdigitale* MR 816 assembly. The Picard tools (http://picard.sourceforge.net ) AddOrReplaceReadGroups, MarkDuplicates, CreateSequenceDictionary, and ReorderSam were used to preprocess read alignments. To minimize false positive SNP calls near insertion/deletion (indel) events, poorly aligned regions were identified and realigned using GATK RealignerTargetCreator and IndelRealigner (GATK version 2.7-4 [41]). SNPs were identified using the GATK UnifiedGenotyper (with the haploid genotype likelihood model) run with the SNP genotype likelihood models (GLM). We also ran BaseRecalibrator and PrintReads for base quality score recalibration on sites called using GLM SNP and re-called variants with UnifiedGenotyper emitting all sites. VCFtools [42] was used to count SNP frequency in windows across the genome (--SNPdensity 5000) and to measure nucleotide diversity (--site-pi), which was normalized for the assembly size. For comparison, the nucleotide diversity was calculated for the SNPs identified in a set of 159 isolates of *C. neoformans* var. *grubii*, a fungal pathogen that undergoes frequent recombination [43].

SNPs were mapped to genes using VCFannotator (http://vcfannotator.sourceforge.net/), which annotates whether a SNP results in synonymous or non-synonymous change in coding region. The total number of synonymous and non-synonymous sites across the *T. rubrum* CBS 118829 and *T. interdigitale* MR 816 gene sets were calculated across all coding regions using codeml in PAML (version 4.8) [44]; these totals were used to normalize the ratios of non-synonymous to synonymous SNPs.

### Copy number variation

To identify regions of *T. rubrum* that exhibit copy number variation between the isolates, we identified windows showing significant variation in normalized read depth using CNVnator [45]. The realigned read files used for SNP calling were input to CNVnator version 0.2.5, specifying a window size of 1kb. Regions reported as deletions or duplications were filtered requiring p-val1 < 0.01.

### Phylogenetic and comparative genomic analysis

To infer the phylogenetic relationship of the sequenced isolates, we identified single copy genes present in all genomes using OrthoMCL [46]. Individual orthologs were aligned with MUSCLE [47] and then the alignments were concatenated and input to RAxML [48], version 7.3.3 with 1,000 bootstrap replicates and model GTRCAT. RAxML version 7.7.8 was used for phylogenetic analysis of SNP variant in seven *T. rubrum* isolates, with the same GTRCAT model.

For each gene set, HMMER3 [49] was used to identify PFAM domains using release 27 [50]; significant differences in gene counts for each domain were identified using Fisher’s exact test, with p-values corrected for multiple testing [51]. Proteins with LysM domains were identified using a revised HMM as previously described [15]; this HMM includes conserved features of fungal LysM domains including conserved cysteine residues not represented in the PFAM HMM model and identified additional genes with this domain.

### Construction of paired allele compatibility matrix

To construct SNP profiles, SNPs shared by at least two members of the *T. rubrum* dataset were selected. Private SNPs are not informative for a paired allele compatibility test because they can never produce a positive result. These profiles were then counted across the genome to construct SNP profiles via a custom Perl script. We required profiles to be present at least twice, to minimize the signal from homoplasic mutations. Pairwise tests were then conducted between each of the profiles to look for all four possible allele combinations, which would only occur via either mating or homoplasic mutations.

### Linkage disequilibrium calculation

Linkage disequilibrium was calculated for *T. rubrum* SNPs in 1kb windows of all scaffolds with VCFtools version 1.14 [42], using the --hap-r2 option with a minimum minor allele frequency of 0.2.

### Data Availability

All genomic data is available in NCBI and can be accessed via the accession numbers in Table S2. The NCBI GenBank accession numbers of the three MLST loci are listed in Table S3.

## Results

### Relationship of global Trichophyton isolates using MLST

To examine the relationship of global isolates of *T. rubrum*, we sequenced three loci in each of 104 *Trichophyton* isolates and carried out phylogenetic analysis. The typed isolates included 91 *T. rubrum* isolates, 11 *T. interdigitale* isolates, and 2 *T. benhamiae* isolates (**Table S1**). In addition, data from the genome assemblies of additional dermatophyte species (*T. verrucosum*, *T. tonsurans*, *T. equinum*, and *M. gypseum*) were also included. Three loci — the *TruMDR1* ABC transporter [19], an intergenic region (IR), and an alpha-1,3-mannosyltransferase (CAP59 protein domain) — were sequenced in each isolate. Phylogenetic analysis of the concatenated loci can resolve species boundaries between the seven species (**Figure 1**). A large branch separates a *T. benhamiae* isolate (MR857) from the previously described genome sequenced isolate (CBS 112371) (**Figure 1**), and the sequences of two loci of a second *T. benhamiae* isolate (MR827) were identical to those of MR857 (**Table S3**). Sequencing of the ITS region of the MR857 and MR827 isolates revealed high sequence similarity to isolates from the *T. benhamiae* African race (**Methods**), which is more closely related to *T. bullosum* than isolates of *T. benhamiae* Americano-European race including CBS112371 [52]. Otherwise, the species relationships and groups are consistent between studies.

**Figure 1.**
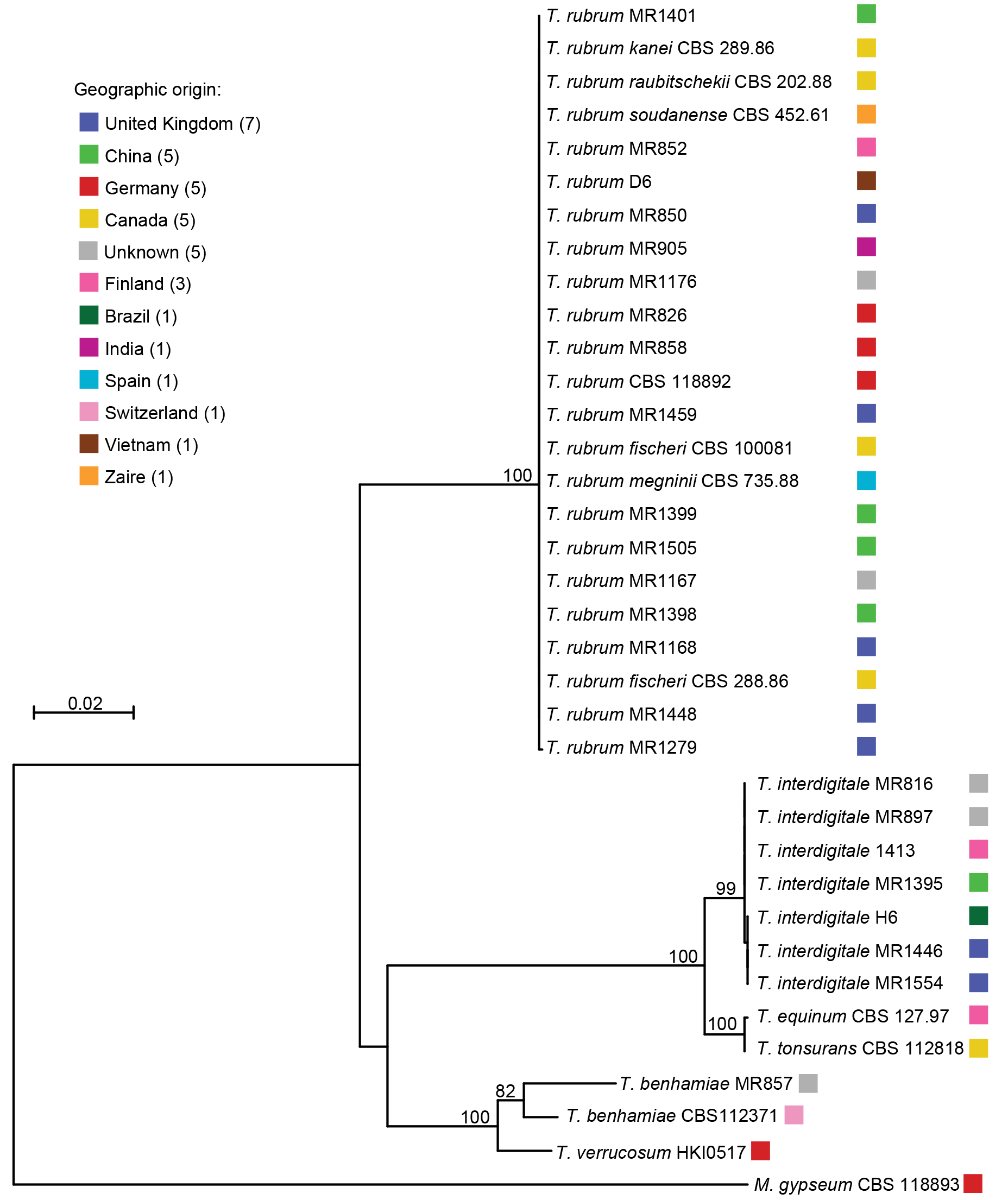
Phylogeny inferred from concatenated MLST sequences. Three MLST loci (ABC transporter, outer membrane protein, and CAP59 protein) were amplified and sequenced from 79 isolates and sequences were identified in an additional 19 assemblies. The concatenated sequence for each isolate was used to build a maximum likelihood tree using MEGA 5.2. Isolate MR1168 is representative of 73 *T. rubrum* isolates that have identical MLST sequences.

MLST analysis demonstrated that the *T. rubrum* isolates were nearly identical at the three sequenced loci. Remarkably, of the 84 *T. rubrum* isolates sequenced at all three loci, 83 were identical at all positions of the three loci sequenced (genotype 2, **Table S3**). Only one isolate, 1279, displayed a single difference at one site in the *TruMDR1* gene (genotype 3, **Table S3**). For the remaining six isolates, sequence at a subset of the loci was generated and matched that of the predominant genotype. Thus, MLST was not sufficient to discern the phylogenetic substructure in the *T. rubrum* population that included six isolates representing different morphotypes (**Table S3**). Similarly, the 11 *T. interdigitale* isolates were highly identical at these three loci; two groups were separated by a single nucleotide difference in the IR and the third group contained a 6-base deletion overlapping the same base of the IR (genotypes 1, 5 and 6, **Table S3**). Although most species can be more easily discriminated based on the MLST sequence, *T. equinum* and *T. tonsurans* isolates differred only by a single transition mutation in the IR, which illustrates the remarkable clonality of these species.

### Genome sequencing and refinement of phylogenetic relationships

As MLST analysis was insufficient to resolve the population substructure of the *T. rubrum* species complex, we sequenced the complete genomes of *T. rubrum* isolates representing worldwide geographical origins and five morphotypes: *fischeri*, *kanei*, *megninii*, *raubitschekii*, and *soudanense*. We generated whole genome Illumina sequences for ten *T. rubrum* and two *T. interdigitale* isolates (**Table S2**). The sequence of each isolate was assembled and utilized to predict gene sets. The *T. rubrum* assembly size was very similar across isolates, ranging from 22.5 to 23.2 Mb (**Table S5**). The total predicted gene numbers were also similar across the isolates, with between 8,616 and 9,064 predicted genes in the ten *T. rubrum* isolates, and 7,993 and 8,116 predicted genes in the two *T. interdigitale* isolates (**Table S5**).

To infer the phylogenetic relationship of these isolates and other previously sequenced *Trichophyton* isolates, we identified 5,236 single-copy orthologs present in all species and estimated a phylogeny with RAxML [48] (**Figure 2A**). This phylogeny more precisely delineates the species groups than that derived from the MLST loci and also illustrates the relationship between the *T. rubrum* isolates (**Figure 2B**). The results of this analysis suggest that the *fischeri* morphotype is not monophyletic, as one *fischeri* isolate (CBS100081) is more closely related to the *raubitschekii* isolate than to the other *fishcheri* isolate (CBS 288.86). While a subset of seven *T. rubrum* isolates appear closely related, others show much higher divergence, including the *soudanense* isolate, the *megninii* isolate, the MR1459 isolate, and the CBS 118829 isolate representing the reference genome. The *soudanense* isolate (CBS 452.61) was placed as an outgroup relative to the other *T. rubrum* isolates; this is consistent with this isolate being part of a clade more closely related to *T. violaceum* than to *T. rubrum* [6] and with the re-establishment of *soudanense* isolates as a separate species [3].

**Figure 2.**
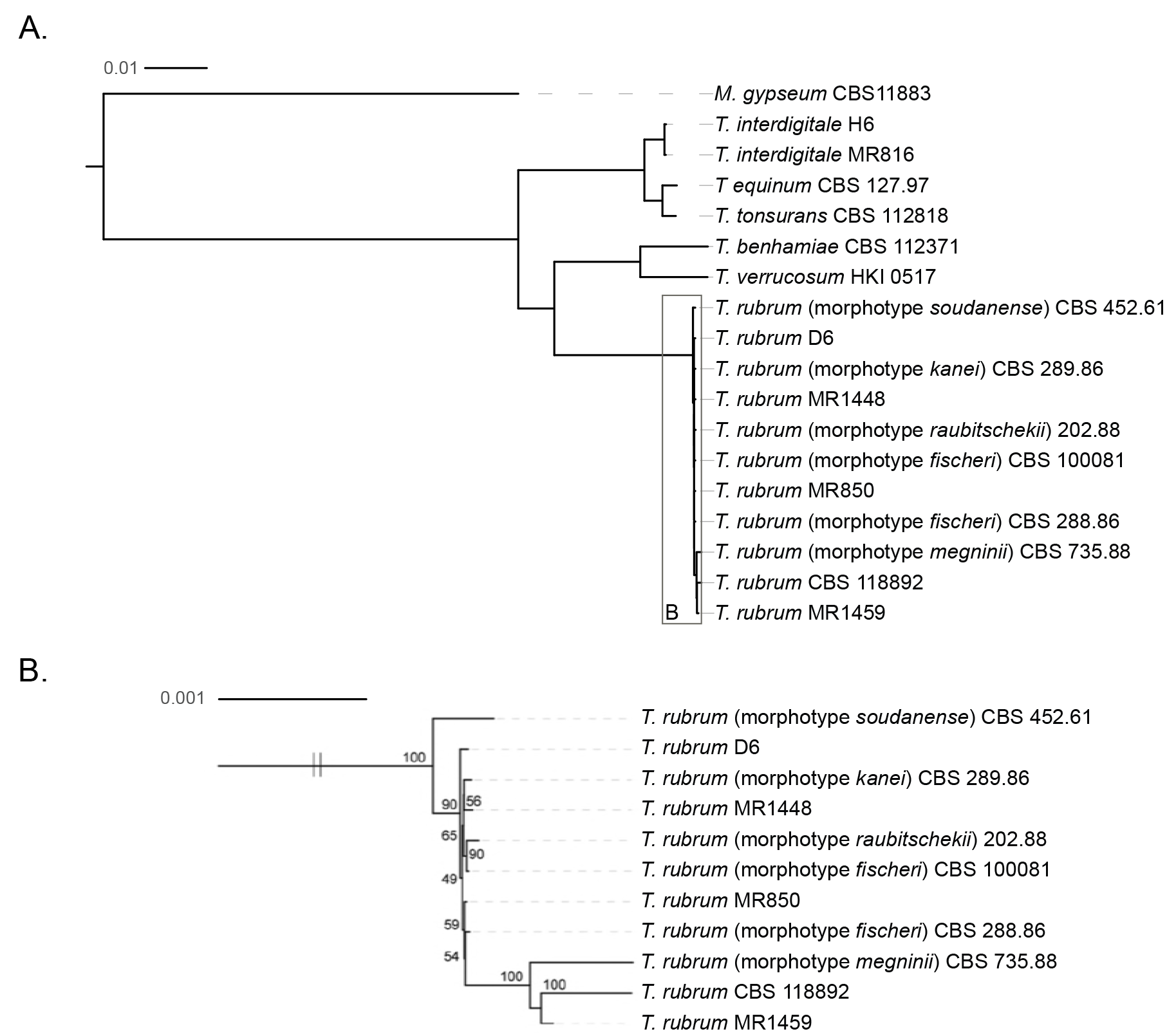
Phylogenetic relationship of *Trichophyton* isolates. A total of 5,236 single copy genes were each aligned with MUSCLE; the concatenated alignment was used to infer a species phylogeny with RAxML (GTRCAT model) with 1,000 bootstrap replicates using either A. all species including the outgroup *M. gypseum* or B. only *Trichophyton rubrum* isolates.

To further classify the two *T. interdigitale* isolates, we assembled the ITS region of the ribosomal DNA locus and compared the sequences to previously classified ITS sequences, as *T. interdigitale* isolates differ from *T. mentagrophytes* at the ITS locus [3,53]. For the two genomes of these species that we sequenced, MR816 was identical to *T. interdigitale* at the ITS1 locus, wheras the H6 isolate appears intermediate between *T. interdigitale* and *T. mentagrophytes*, containing polymorphisms specific to each group (**Figure S1**). Genomic analysis of allele sharing across a wider set of *T. interdigitale* and *T. mentagrophytes* isolates could be used to evaluate the extent of hybrid genotypes and genetic exchange between these two species.

### *MAT1-1* prevalence and clonality in *T. rubrum*

To address if the *T. rubrum* population is capable of sexual reproduction, we surveyed the *MAT* locus of all isolates. Using either gene content in assembled isolates or a PCR assay to assign mating type, we found that 79 of the 80 *T. rubrum* isolates contained the alpha domain gene at the *MAT* locus (*MAT1-1*). In addition, a set of 55 isolates from Turkey were found to harbor the *MAT1-1* allele based on a PCR assay (**Figure S2**). However, the *T. rubrum* morphotype *megninii* isolate contained an HMG gene at the *MAT* locus (*MAT1-2*) (**Figure 3**, **Table S1**). The presence of both mating types suggests that this species could be capable of mating under some conditions. However the high frequency of a single mating type strongly suggests that *T. rubrum* largely undergoes clonal growth, although other interpretations are also possible (see **Discussion**). In further support of this, a study of 206 *T. rubrum* clinical isolates from Japan noted that all were of the *MAT1-1* mating type [14].

**Figure 3.**
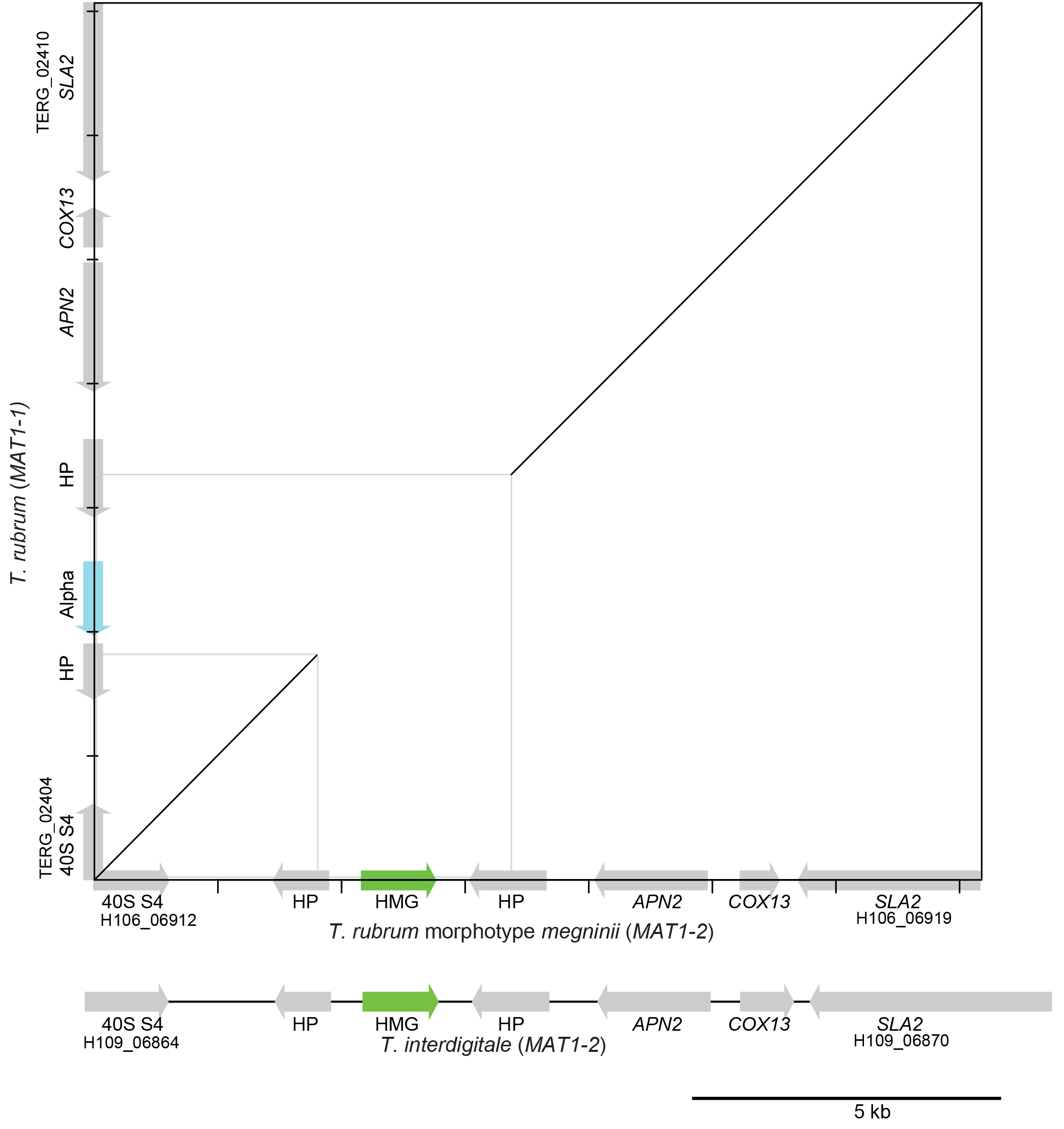
Alignment of the mating type locus of selected isolates. Mating type genes of *T. rubrum* morphotype *megninii* (CBS 735.88) and *T. rubrum* (CBS 188992) are shown along the x- and y-axes, respectively, with regions aligning by NUCmer show in the dotplot. The alignment extends into two hypothetical proteins (HP) immediately flanking the alpha or HMG domain gene that specifies mating type. Most *T. rubrum* (*MAT1-1*) isolates contain an alpha domain protein (blue) at the *MAT* locus. In contrast, the *T. rubrum* morphotype *megninii* isolate contains an HMG domain protein(green) representing the opposite mating type (*MAT1-2*). All sequenced *T. interdigitale* isolates are also of *MAT1-2* mating type including MR816. Gene locus identifiers are shown for the genes flanking each locus (prefix TERG, H106, and H109).

A closer comparison of the genome sequences of *T. rubrum* isolates also supports a clonal relationship of this population. Phylogenetic analysis of the seven most closely related *T. rubrum* isolates using SNPs between these isolates (see below) suggests that the isolates have a similar level of divergence from each other (**Figure S3**). This supports that these *MAT1-1 T.rubrum* isolates have likely undergone clonal expansion.

To test for recombination that could reflect sexual reproduction within the *T. rubrum* population sampled here, we conducted a genome-wide paired allele compatibility test to look for the presence of all four products of meiosis (**Figure 4**). This test is a comparison between two paired polymorphic sites in the population. While the presence of three of the four possible allele combinations at two sites in a population is possible through a single mutation and identity by descent, the presence of all four combinations requires either recombination, or less parsimoniously, a second homoplasic mutation. Four positive tests resulted from this analysis (out of 21 possible), including allele combinations that occurred a minimum of 13 times. This may suggest that recombination is a rare event arising through infrequent sexual recombination occurring in this population although the same mutations and combinations arising via homoplasy (or selection) are difficult to exclude. Based on the number of triallelic sites in the dataset (19), we would predict 9.5 homoplasic sites to have occurred by random chance, which is similar to the number of sites responsible for the positive signals in the compatibility test. In addition, linkage disequilibrium does not decay over increasing distance between SNPs in *T. rubrum* (**Figure S4**), which further supports a low level of recombination in this species; sequencing additional diverse isolates would help to address if some isolates or lineages were more prone to recombination.

**Figure 4.**
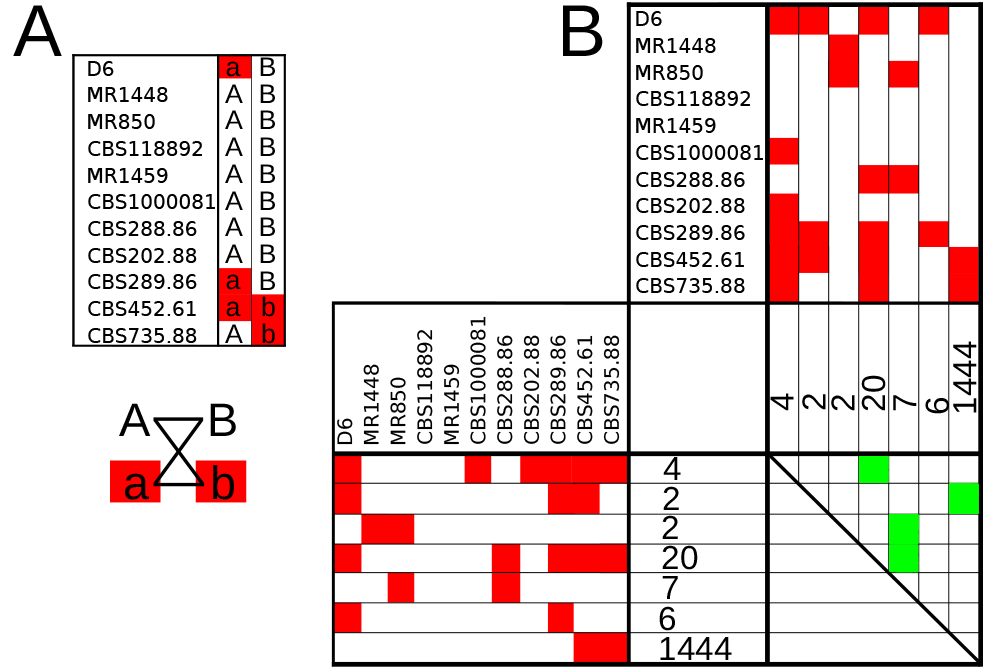
Paired allele compatibility test suggests limited evidence for sexual reproduction. A. A single example of a positive paired allele compatibility test from the *T. rubrum* population. In this test, two loci are examined and typed across the population. To perform a meaningful test, at least two individuals in the population must share a variant allele at each site. Here alternative SNPs are depicted in red and reference in white. Evidence for recombination is provided by any pairwise comparison of two loci in which isolates are present where red-red, white-white, red-white, and white-red combinations are all found (AB, Ab, aB, and ab) satisfying the allele compatibility test and providing evidence for recombination. B. Paired allele compatibility tests were performed for all isolates in the *T. rubrum* population across the entire genome. SNP profiles were grouped into unique and informative allele patterns and collapsed, with the number of occurrences of each profile across the genome listed. Thus, the larger the number, the more common that SNP distribution is in the population. Pairwise tests were then conducted for each combination of SNP profiles. Reference nucleotides are indicated by white and variant by red. The pairwise matrix displays the results of all of these tests; a green square in the pairwise matrix is indicative of a positive test for the pairwise comparison and thus provides potential evidence of recombination.

We also characterized the *MAT* locus of the newly sequenced *T. interdigitale* isolates (H6 and MR816) and found that both contain an HMG domain gene. These *T. interdigitale* isolates were more closely related to *T. equinum* (*MAT1-2*) and *T. tonsurans* (*MAT1-1*) than *T. rubrum* (**Figure 2A**). To survey the mating type across a larger set of *T. interdigitale* isolates, a set of 11 additional isolates from Turkey were typed. Based on PCR analysis, all *T. interdigitale* isolates harbor the *MAT1-2* allele (**Figure S2**).

The mating abilities of the isolates were tested by conducting mating assays with potentially compatible isolates of *T. rubrum*, including the *megninii* morphotype, *T. interdigitale*, and *T. simii* (**Table S7**). These experiments were conducted using both Takashio and E medium at room temperature (approximately 21 to 22°C) without Parafilm in the dark. Although the assay plates were incubated for longer than five months, ascomata or ascomatal initials were not observed (**Figure S5**). While it is possible that mating may occur under cryptic conditions [54], this data suggests that the conditions tested are not sufficient for the initiation of mating structures in *T. rubrum*.

### Genome-wide variation patterns in *T. rubrum*

SNP variants were identified between *T. rubrum* isolates to examine the level of divergence within this species complex (**Table S8**). On average, *T. rubrum* isolates contain 8,092 SNPs compared to the reference genome of the CBS118892 isolate; this reflects a bimodal divergence pattern where most isolates including three morphotypes (*fischeri*, *kanei*, and *raubitscheckii*) have an average of 3,930 SNPs and two more divergent isolates (morphotypes *megninii* and *soudanense*) have an average of 24,740 SNPs. The average nucleotide diversity (TT) for all 10 *T. rubrum* isolates is 0.00054; excluding the two divergent morphotypes, the average nucleotide diversity is 0.00031. By comparison, the average nucleotide diversity of the fungal pathogen *Cryptococcus neoformans* var. *grubii*, which is actively recombining as evidenced by low linkage disequilibrium [43,55], is 0.0074, a level approximately 24-fold higher than that in *T. rubrum* (**Methods**, [43]). Even higher levels of nucleotide diversity have been reported in global populations of other fungi (see **Discussion**). A similar magnitude of SNPs separate the two *T. interdigitale* isolates; 22,568 SNPs were identified based on the alignment of H6 reads to the MR 816 assembly. Across all isolates, SNPs were predominantly found in intergenic regions for both species, representing 76% and 81% of total variants respectively (**Table 1**, **Table S8**). Within genes, the higher ratio of nonsynonymous relative to synonymous changes among the closely related *T. rubrum* isolates (**Table 1**) is consistent with lower purifying selection over recent evolutionary time [56].

**Table 1.** Variation in T. rubrum SNP rate and class

| Isolate | Total Number of SNPs | SNPs in CDS* | SYN* | NSY* | pN/pS* |
| --- | --- | --- | --- | --- | --- |
| *T. rubrum* MR1448 | 4,283 | 374 | 83 | 287 | 1.15 |
| *T. rubrum* MR1459 | 2,188 | 436 | 103 | 317 | 1.02 |
| *T. rubrum* MR850 | 4,203 | 387 | 88 | 289 | 1.09 |
| *T. rubrum* D6 | 4,121 | 484 | 112 | 363 | 1.08 |
| *T. rubrum* (morphotype *fischeri*) CBS 100081 | 4,199 | 409 | 94 | 307 | 1.09 |
| *T. rubrum* (morphotype *fischeri*) CBS 288.86 | 4,147 | 375 | 84 | 283 | 1.12 |
| *T. rubrum* (morphotype *kanei*) CBS 289.86 | 4,491 | 474 | 116 | 350 | 1.00 |
| *T. rubrum* (morphotype *raubitschekii*) CBS 202.88 | 3,808 | 375 | 83 | 285 | 1.14 |
| *T. rubrum* (morphotype *megninii*) CBS 735.88 | 26,406 | 7,328 | 3,069 | 4,185 | 0.45 |
| *T. rubrum* (morphotype *soudanense*) CBS 452.61 | 23,073 | 6,253 | 2,377 | 3,808 | 0.53 |
| *T. interdigitale* MR816 | 1,223,298 | 591,173 | 395,250 | 194,498 | 0.16 |
| *T. interdigitale* H6 | 1,183,411 | 585,288 | 393,079 | 190,826 | 0.16 |

\*CDS, coding sequence; SYN, synonymous SNP sites; NSY, non-synonymous SNP sites; pN/pS, (NSY/total NSY sites)/(SYN/total SYN sites).

Examining the frequency of SNPs across the *T. rubrum* genome revealed high diversity regions that flank the mating type locus in the two divergent isolates. Across all isolates, some regions of the genome are over-represented for SNPs, including the smallest scaffolds of the reference genome (**Figure 5**); these regions contain a high fraction of repetitive elements [15]. The largest high diversity window unique to the *T. rubrum* morphotype *megninii* was found in an ∼810-kb region encompassing the mating type locus on scaffold 2; a smaller high diversity region spanning the mating type locus was found in the diverged *soudandense* isolate (**Figure 5**). The higher diversity found in this location could reflect introgressed regions from recent outcrossing or could be associated with lower recombination proximal to the mating type locus, resulting in stratification of linked genes.

**Figure 5.**
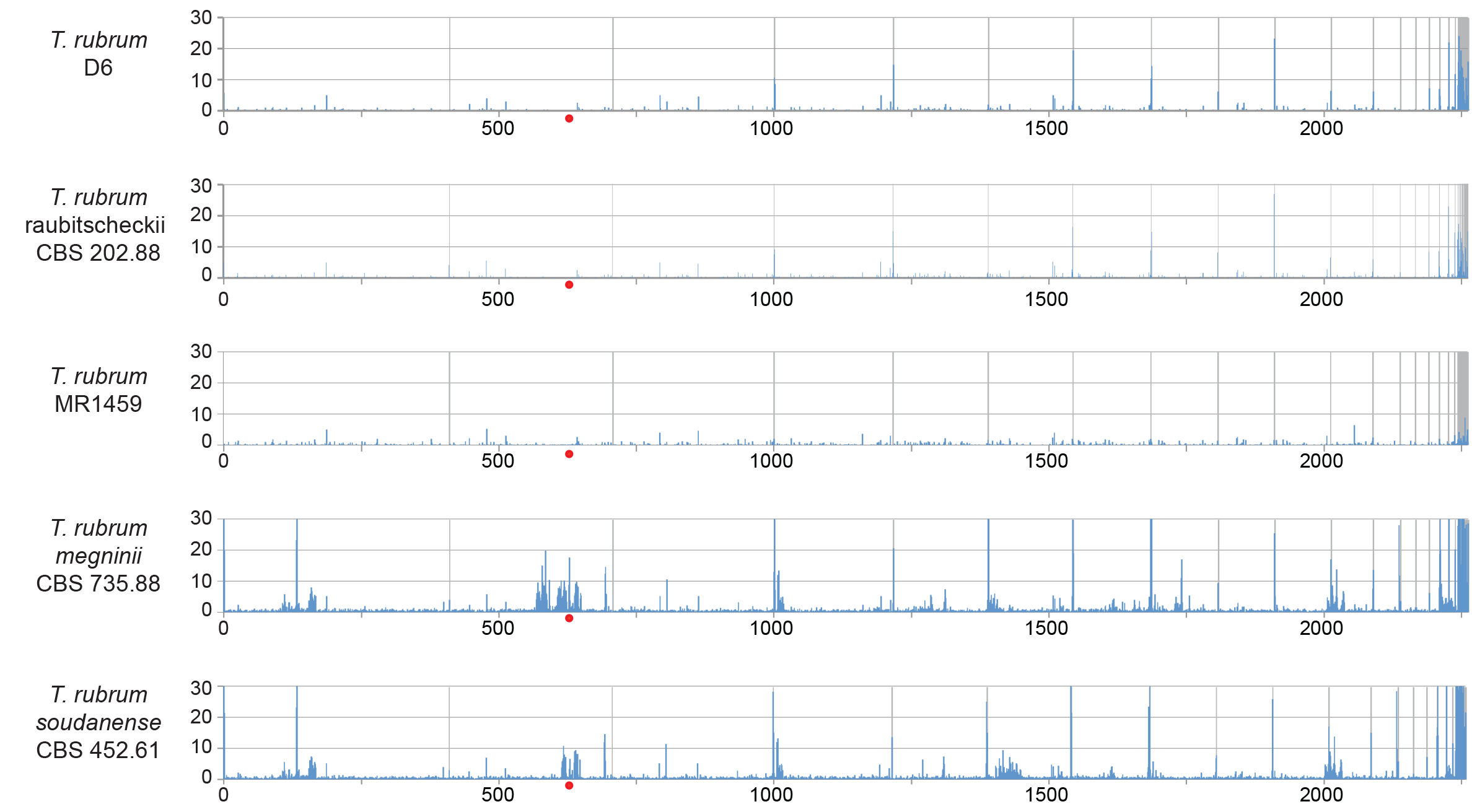
Genome-wide SNP frequency highlights hotspots. For each panel, the frequency of SNPs in 5-kb windows is shown across the genome. The genome assembly of isolate CBS 11892 was used for all comparisons, and scaffolds are ordered along the x-axis with grey lines representing scaffold boundaries. Red dots indicate the position of the mating type locus.

### Gene content variation in *T. rubrum* and *T. interdigitale*

To examine variation in gene content in the *Trichophyton rubrum* species complex, we first measured copy number variation across the genome. Duplicated and deleted regions of the genome were identified based on significant variation in normalized read depth (Methods). We observed increased copy number only for two adjacent 26 kb regions of scaffold 4 in two isolates (MR850 and MR1448) (**Figure S6**). Both of these regions had nearly triploid levels of coverage (**Table S9**). While ploidy variation is a mechanism of drug resistance in fungal pathogens, none of the 25 total genes in these regions (**Table S10**) are known drug targets or efflux pumps. These regions include two genes classified as fungal zinc cluster transcription factors; this family of transcription factors was previously noted to vary in number between dermatophyte species [15]. A total of 12 deleted regions (CNVnator p-val <0.01) ranging in size from 4 to 37 kb were also identified in a subset of genomes (**Table S11**). Two of these regions include genes previously noted to have higher copy number in dermatophyte genomes, a nonribosomal peptide synthase (NRPS) gene (TERG_02711) and a LysM gene (TERG_02813) (**Table S12**). Overall this analysis suggests recent gain or loss in dermatophytes for a small set of genes including transcription factors, NRPS, and LysM domain proteins.

We next examined candidate loss of function mutations in the *T. rubrum* species complex. For the 8 closely related *T. rubrum* isolates, an average of 8.1 SNPs are predicted to result in new stop codons, disrupting protein coding regions; in the *soudanense* and *megninii* isolates, an average of 58.5 SNPs result in new stop codons. These predicted loss of function mutations do not account for previously noted phenotypic differences between the morphotypes; no stop codons were found in the seven genes involved in histidine biosynthesis (*HIS1*-*HIS7*) in the histidine auxotroph *T. rubrum* morphotype *megninii* or in urease genes in *T. rubrum* morphotype *raubitscheckii*.

Comparison of the first representative genomes for *T. interdigitale* (isolates MR816 and H6) to those of dermatophyte species highlighted the close relationship of *T. interdigitale* to *T. tonsurans* and *T. equinum*. These three species are closely related (**Figure 2**), sharing 7,618 ortholog groups, yet there are also substantial differences in gene content. A total of 1,253 orthologs groups were present only in *T. equinum* and *T. tonsurans* and 512 ortholog groups were present only in both *T. interdigitale* isolates. However, there were no significant differences in functional groups between these species based on PFAM domain analysis, suggesting no substantial gain or loss of specific protein families. Two PFAM domains were unique to the *T. interdigitale* isolates and present in more than one copy: PF00208, found in ELFV dehydrogenase family members and PF00187, a chitin recognition protein domain. This chitin binding domain is completely absent from the *T. equinum* and *T. tonsurans* genomes while in *T. interdigitale* this domain is associated with the glycosyl hydrolase family 18 (GH18) domain [57]. GH18 proteins are chitinases and some other members of this family also contain LysM domains. We also examined genes in the ergosterol pathway for variation, as this could relate to drug resistance; while this pathway is highly conserved in dermatophytes [15], *T. interdigitale* isolates had an extra copy of a gene containing the ERG4/ERG24 domain found in sterol reductase enzymes in the ergosterol biosynthesis pathway. The *ERG4* gene encodes an enzyme that catalyzes the final step in ergosterol biosynthesis, and it is possible that an additional copy of this gene results in higher protein levels to help ensure that this step is not rate limiting.

These comparisons also highlighted the recent dynamics of the LysM family, which binds bacterial peptidoglycan and fungal chitin [58]. Dermatophytes contain high numbers of LysM domain proteins ranging from the 10 genes found in *T. verrucosum* to 31 copies found in *M. canis* (**Table S13, [15]**). Both the class of LysM proteins with additional catalytic domains and the larger class consists of proteins with only LysM domains, many of which contain secretion signals and may represent candidate effectors [15], vary in number across the dermatophytes. Isolates from the *T. rubrum* species complex have 16 to 18 copies of LysM proteins compared to the 15 found in the previously reported genome of the CBS 118892 isolate (**Table S13**). One of the additional LysM genes present in all of the newly sequenced isolates encodes a polysaccharide deacetylase domain involved in chitin catabolism. There is also an additional copy of a gene with only a LysM domain in 9 of the 10 new *T. rubrum* isolates (**Table S13**). The genomes of the *T. interdigitale* isolates have only 14 genes containing a LysM binding domain, and are missing a LysM gene encoding GH18 and Hce2 domains (**Figure S7**). Notably, this locus is closely linked to genes encoding additional LysM domain proteins in some species (**Figure S7**). The variation observed in the LysM gene family suggests that recognition of chitin appears to be highly dynamic based on these differences in gene content and domain composition.

## Discussion

In this study, we selected diverse *T. rubrum* isolates for genome sequencing, assembly, and analysis and surveyed a wider population sample using MLST analysis. These isolates include multiple morphotypes, which show noted phenotypic variation yet are assigned to the same species based on phylogenetic analyses [3,53]. The *T. rubrum* morphotype *soudanense* and *T. rubrum* morphotype *megninii* show higher divergence from a closely related subgroup that includes the *kanei*, *raubitschekii*, and *fischerii* morphotypes, as well as most other *T. rubrum* isolates.

Our MLST and whole genome analyses provide strong support that *T. rubrum* is highly clonal and may be primarily asexual or at least infrequently sexually reproducing. Across 135 isolates examined, 134 were from a single mating type (*MAT1-1*). Only the *T. rubrum* type *megninii* isolate, Consistent with prior reports [3,53], only the *T. rubrum* morphotype *megninii* isolates contains the opposite mating type (*MAT1-2*) while all other *T. rubrum* isolates that are of *MAT1-1* type. Direct tests of mating between these and other species did not find evidence for mating and sexual development. While mating was not detected, studies in other fungi have required specialized conditions and long periods of time to detect sexual reproduction [59]. As genes involved in mating and meiosis are conserved in *T. rubrum* [15], gene loss does not provide a simple explanation for the inability to mate. Sexual reproduction might occur rarely under specific conditions such as specific temperatures as found for *Trichophyton onychocola* [60], may be geographically restricted, as the opposite mating type *megninii* morphotype is generally found in the Mediterranean [61], or could be unisexual as in some other fungi such as *Cryptococcus neoformans* [62].

As MLST data provided no resolution of the substructure of the *T. rubrum* population, we examined whole genome sequences for 8 diverse isolates. Analysis of the sequence read depth revealed that while some small regions of the genome show amplification or loss, there is no evidence for aneuploidy of entire chromosomes. Most of these *T. rubrum* isolates contain an average of only 3,930 SNPs (0.01% of the genome) and phylogenetic analysis revealed little genetic substructure. Two isolates were more divergent with an average of 24,740 SNPs (0.06% of the genome); one of these was of the recently proposed separated species *T. soudanense* [3], and the other was the *T. rubrum* morphotype *megninii* isolate. While the similar level of divergence raises the question of whether morphotype *megninii* isolates could also be a separate species, this has not yet been proposed when considering additional phenotypic data in addition to molecular data, however further study would help clarify species assignments. The low level of variation is remarkable in comparison to other fungal pathogens; for example, while *T. rubrum* isolates are identical at 99.97% of positions on average, isolates of *Cryptococcus neoformans* var. *grubii* isolates are 99.36% identical on average [43,55]. Global populations of *Saccharomyces* have even higher reported diversity [63]. The low diversity and the dependency on the human host for growth suggests that *T. rubrum* may have a low effective population size impacted by the reduction of intra-species variation by genetic drift. In addition, direct tests for recombination found a low level of candidate reassortments that was not in excess of the estimated number of homoplasmic mutations; further, as there was no apparent decay of linkage disequilibrium over genetic distance, our analyses support the overall clonal nature of this species. The high clonality observed in *T. rubrum* is also supported by MLST analysis of eight microsatellite markers in approximately 230 *T. rubrum* isolates, including morphotypes from diverse geographic origins [8]. With additional genome sequencing geographic substructure may become more apparent; the fungal pathogen *Talaromyces marneffei* also displays high clonality yet isolates from the same country or region are more closely related [64]. While low levels of diversity seems surprising in a common pathogen, this is similar to findings in some bacterial pathogens including *Mycobacterium tuberculosis* and *Mycobacterium leprae* [65,66], which also display high clonality despite phenotypic variation.

LysM domain proteins are involved in dampening host recognition of fungal chitin [67] and can also regulate fungal growth and development [68], yet their specific function in dermatophytes and closely related fungi is not well understood. We also observed variation in genes containing the LysM domain across the sequenced isolates, both in the gene number and domain organization. LysM genes are present in higher copy number in dermatophytes than related fungi in the Ascomycete order Onygenales [15]. Recent sequencing of additional non-pathogenic species in this order related to *Coccidioides* revealed that most LysM copies found in dermatophytes have a homolog [69]. Although this analysis excluded *M. canis* — the dermatophyte species with the highest LysM count— this suggests that dermatophytes have retained rather than recently duplicated many of their LysM genes. However, changes in the domain composition of both genes with catalytic domains and those with only LysM domains, many of which represent candidate effectors, highlights the dynamic evolution of the LysM family in the dermatophytes. Studies of LysM genes in dermatophytes are needed to determine whether these genes serve similar or different roles in these species.

*T. rubrum* is only found as a pathogen of humans, though this adaptation is more recent relative to the related species that infect other animals or grow in the environment. Unlike the obligate human fungal pathogen *Pneumocystis jirovecii* [70,71], *T. rubrum* does not display widespread gene loss [15] indicative of host dependency for growth; further, its genome size is also comparable to related dermatophyte species, supporting no overall reduction [15]. The presence of a single mating type in the vast majority of isolates and limited evidence of recombination suggests that sexual reproduction of *T. rubrum* may have been recently lost or may be rarely occurring in specific conditions or geographic regions. This may be linked to the specialization as a human pathogen, as mating may be optimized during environmental growth in the soil [53].

## Acknowledgements

We thank the Broad Institute Genomics Platform for generating the DNA sequence described here. We thank Yonathan Lewit for technical assistance and Cecelia Wall for providing helpful comments on the manuscript. Financial support was provided by the National Human Genome Research Institute grant number U54HG003067 to the Broad Institute and by NIH/NIAID R37 MERIT Award AI39115-20 and RO1 Award AI50113-13 to JH. This study was supported by The Scientific and Technological Research Council of Turkey-2219 Research Fellowship Programme for International Researchers Project No. [1059B191501539] to AD and Brazilian funding agency FAPESP Fundação de Amparo à Pesquisa do Estado de São Paulo, Postdoctoral Fellowship 12/22232-8 and 13/19195-6 to GFP.

## Author contributions

C.A.C., D.A.M., T.C.W. and J.H. conceived and designed the project. A.D., B.M, S.H., M. I., R.B., B.O., Y.G, N.M.M, and T.W. provided the isolates. W.L. and A.D. performed the laboratory experiments. G.F.P, D.A.M, W.L, A.D., R.B.B, A.A, J.M., G,T.S., S.Y, Q.Z, and C.A.C analyzed the data. C.A.C. and J.H. wrote the paper with input from all authors. C.A.C. and J.H. supervised and coordinated the project.

## Supplemental Figure and Table Legends

**Figure S1. ITS sequence variation in *T. interdigitale***. Aligned ITS sequence is shown for four isolates, including the two for which whole genomes were sequenced (H6 and MR816) and two previously characterized isolates (AF168124 and AY062119). Isolate AY062119 has been re-classified as *T. mentagrophytes*. Variant sites are highlighted.

**Figure S2. Detection of *MAT1-1* in *T. rubrum* and *MAT1-2* in *T. interdigitale* by PCR**. A. PCR-based determination of the *MAT1-1* alpha domain of *T. rubrum* isolates: *T. rubrum* MR 851, *T*. *megninii* CBS 389.58, *T. megninii* CBS 384.64, and *T*. *megninii* CBS 417.52. The alpha domain *MAT1-1* gene was identified from samples = 1-12 and MR 851 *T. rubrum*; the alpha domain *MAT1-1* gene was not identified from *T*. *megninii* CBS 389.58, *T*. *megninii* CBS 384.64, or *T*. *megninii* CBS 417.52. M: DNA ladder. B. PCR-based determination of *MAT 1-2* HMG domain of *T. rubrum*: *T. rubrum* MR851, *T*. *megninii* CBS 389.58, *T*. *megninii* CBS 384.64, and *T*. *megninii* CBS 417.52. The HMG domain *MAT1-2* gene was identified from *T*. *megninii* CBS 389.58, *T*. *megninii* CBS 384.64, and *T*. *megninii* CBS 417.52, and was not identified from *T. rubrum* isolate = 1-12 and *T* . *rubrum* MR 851, M: DNA ladder.C. PCR-based determination of *MAT1-2* HMG domain of *T. interdigitale. T. interdigitale* isolates = 1-9, *T. interdigitale* MR8801 (positive control), and *T. rubrum* MR851 (negative control), M; DNA ladder. D. PCR based determination of *MAT 1-1* alpha domain of *T. interdigitale* isolates. *T. interdigitale* isolates = 1-9, *T. interdigitale* MR8801 (negative control), and *T. rubrum* MR851 (positive control). M: DNA ladder.

**Figure S3. Phylogenetic relationship and sharing of variant sites of sequenced *T. rubrum* isolates**. A. Phylogenetic relationship of *T. rubrum* isolates inferred using RAxML (Methods). B. Classification of SNP sites based on conservation across the sequenced isolates; unique: only in one isolate; shared: in two to seven isolates; fixed: in all eight isolates.

**Figure S4. Lack of decay of linkage disequilibrium (LD) in *T. rubrum***. LD (r^2^) was calculated for all pairs of SNPs separated by 0 to 300 kb and then averaged for every 1kb. LD values for each window were then calculated by averaging over all pairwise calculations in the window.

**Figure S5. Mating assays**. A. Mating assay plate; *T. rubrum* and *T. interdigitale* on E medium. B. Mating assay plate*; T. rubrum* and *A. simii* (a) E medium, (b) Takashio medium eight weeks. C*. T. rubrum* and *T. megninii* on E medium for eight weeks.

**Figure S6. Read depth of sequenced isolates**. Reads from each isolate were aligned to the *T. rubrum* reference genome and normalized read depth was computed for 5kb windows. Read depth is even across the reference genome for most isolates, with small regions of higher depth detected in some isolates.

**Figure S7. Variation in LysM-Hce gene cluster across sequenced dermatophytes**. In *T. rubrum*, the LysM-Hce gene is closely linked to two other LysM genes; this organization is most similar to that found in *M. canis*, although these genes are located on two different scaffolds.

**Table S1. Properties of sequenced isolates.**

**Table S2. Accessions for sequenced genomes.**

**Table S3. MLST sequence, genotypes, and GenBank accession numbers. Table S4. Primers for MLST gene amplification.**

**Table S5. *Trichophyton* genome assembly statistics.**

**Table S6. Primer pairs used for mating type determination of *T. rubrum***

**Table S7. Mating assays and results.**

**Table S8. Frequency of SNPs in *T. rubrum* and *T. interdigitale* isolates by mutation type.**

**Table S9. Duplicated regions in sequence isolates.**

**Table S10. List of genes found in duplicated regions. Table S11. Deleted regions in sequenced isolates.**

**Table S12. List of genes in deleted regions in sequenced isolate**

**Table S13. Genes containing the LysM binding domain in dermatophytes**

